# Automatically producing large morphometric datasets from natural history collection images: a case study of Lepidoptera wing shape

**DOI:** 10.1101/2022.07.01.497900

**Authors:** Samuel Ginot, Vincent Debat

## Abstract

Publicly available image data (2D and 3D) from biological specimens is becoming extremely widespread, notably following digitization efforts from natural history institutions worldwide. To deal with this huge amount of data, high-throughput phenotyping methods are being developed by researchers, to extract biologically meaningful data, in correlation with the burgeoning of the field of phenomics. Here we explore the potential of a combination of simple image treatment algorithms, with a geometric morphometrics contour analysis, applicable to strongly standardized images such as collections of Lepidotera. Using a previously manually landmarked dataset of *Morpho* butterflies, we show that our automated approach can produce a morphospace similar to that produced by a manual approach. Although the former is more noisy than the latter, it appears to pick up phylogenetic and to some extent ecological signal. Applying then the same approach to a large dataset of images from two different museums, we produce a morphospace containing >5000 specimens, representing 851 species in 24 families of butterflies and moths. The most notable feature of this space is that Sphingidae morphology is clearly separate from the rest, and appears much more constrained. We also show some indirect evidence that at this large interspecific level, potential museum related bias (e.g. inter-user bias in specimen preparation and photography) can be negligible. Altogether, our results suggest that this approach has the potential to produce large-scale analysis of morphology, and could be refined to include more specimens.

## 1 Introduction

Semi-automated image treatment and automated morphometrics can be combined to produce large datasets (Wilson et al., 2022), particularly useful in groups with very numerous specimens and species and in phenomics approaches. Several approaches exist : automatization of landmarking, notably using machine learning, or pattern recognition using neural networks for example (Porto and Voje, 2020; Porto et al., 2021; Lürig, 2022). Recently, one common approach to better quantify morphology has been to increase the number of landmarks to more fully represent these biological shapes, notably using 3D surface semi-landmarks and / or 2D contour (Goswami et al., 2019). However, one remaining challenge is that the number of specimens is technically harder to increase than the number of variables i.e. landmark coordinates), especially for 3D objects, which need to be *μ*CT-scanned, a long and expensive process, but also for 2D objects which need photographed in a standardized fashion to avoid increasing errors too much. One major problem with this is that if the number of variables P » number of individuals N, there tends to be overfitting, for example falsely high rates of correct identifications in discriminant analyses, or apparent patterns that do not hold statistically (Houle et al., 2010; Cardini et al., 2019). To access the phenomics realm, one solution could be to increase sample sizes even more than the number of variables (Houle et al., 2010; Cardini et al., 2019), which, in the case of morphology, could be done using publicly available digital material, such as digital museum collections (3D scans repositories are becoming numerous, while many museums are systematically photographing material). But mixing up sources of data can bring more error noise or biases), and it is not yet sure that automated morphometric methods can be easily applied to such material. The validity of such approaches therefore depends on whether biological signal can be disentangled from non-biological signal (e.g. inter-user differences in preparation and / or digitization of specimens), which depends not only on standardisation, but also on the respective effect sizes of various components in terms of variance. Basically, it comes down to: can having very large sample sizes compensate for the introduced error by making the biological signal more robust? Here we try to answer this question in photographic collections of pinned butterflies, an adequate context because pictures are taken in a standardized fashion even across different institutions: camera directly above the specimen, which lays flat with wings opened and the border separating hind and forewing perpendicular to the major axis of the body. Furthermore, images are generally well contrasted, which allows simple image treatment to separate the specimen from its background. Wings of butterflies are obviously involved in flight and in some cases hover, and the diversity of wing shapes therefore relates to flight type and performance (Le Roy et al., 2019). Using a dataset derived from museum collections we aim to describe butterfly shape variation and try to find out whether it is driven purely by noise or biases, or if taxonomic, phylogenetic and eco-morpho-functional signal can also be detected.

## 2 Material and Methods

### 2.1 Image data

Our sampling is based on pictures of Lepidopteran collections from two natural history museums: the Natural History Museum of London (NHM) and Museum d’Histoire Naturelle de Toulouse (MHNT). All images are publicly available, and can be used under a CC BY-SA licence. Images from the MHNT were digitized by D. Descouens under project Phoebus, and are now also available through the Wikimedia Commons (e.g. Fig. 1). Three image datasets from the NHM were used: the *Birdwing Butterfly Collection* (Wing et al., 2019), the *Papilionoidea New Types Digitisation Project* (Crowther et al., 2019a), and the *iCollections*: *British and Irish Pyraloidea (Moths) Collection* Crowther et al. (2019b).

**Figure 1:**
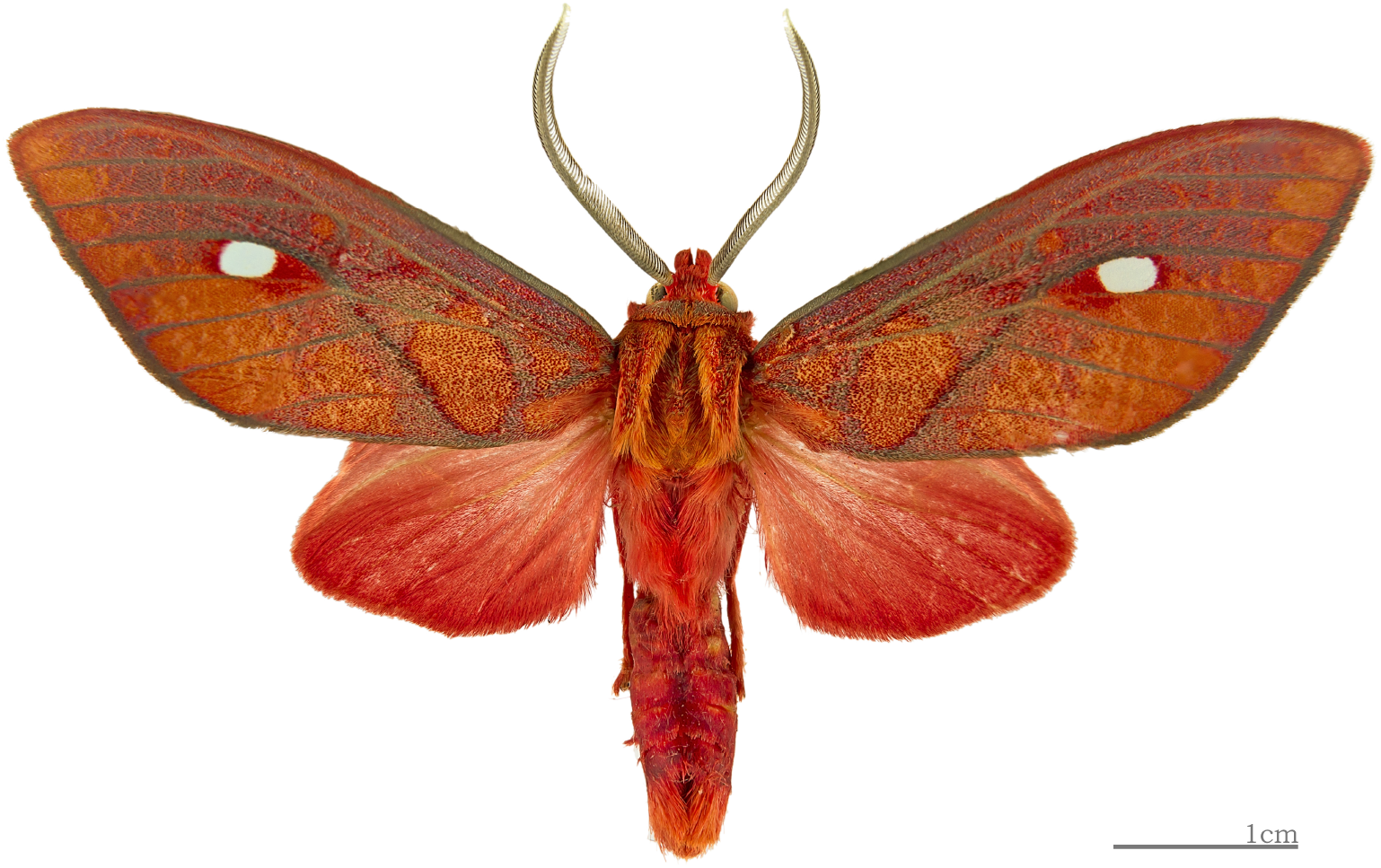
Dorsal view of pinned specimens of a male *Ernassa sanguinolenta* from the MHNT collection. Picture by D. Descouens, MHNT, CC BY-SA.

### 2.2 Automated morphometric data extraction

The following pipeline was applied to the different datasets:

- Get public data from museum collections: Often CSV or TXT files containing information about each specimen, and most importantly their URLs.
- Make text file containing only URLs.
- Automatically download files (Fig. 1), for example using Bash under Linux (caution: tens of GB may be dowloaded, so make sure to have enough space). Under Linux distributions this can be achieved using for example the ‘wget’command.
- Separate the various objects present in the image. Here, we used the *ImageMagick* free command line software (https://imagemagick.org/index.php), with addition of the following scripts from http://www.fmwconcepts.com/imagemagick/index.php: corners, multicrop, multicrop2. Objects that are too small to be of interest are automatically deleted at this step.
- Convert all images to grayscale. Can also be done automatically with *ImageMagick*.
- Manually check that images are correctly cropped and grayscaled.
- From grayscale images, contours of objects are automatically extracted using a custom built R function based on packages ‘EBImage’ (Pau et al., 2010) and ‘Momocs’ (Bonhomme et al., 2014). This function consists of several image treatment actions:
  – Read the image in memory.
  – Translate the resulting object.
  – Floodfill background with black (tolerance parameter has to be adjusted). Since a universal tolerance level can hardly be achieved for thousands of different images, this step may cause some loss in the dataset.
  – Convert grayscale to B / W.
  – Small residual pixels and other objects may still be present in the image if bounding box caught them, so remove them by deleting all objects except background and largest object (which should be the biological item).
  – Invert to black object on white background (Fig. 2).
  – Extract the matrix with pixel values to pass it into the Momocs functions.
  – Auto-extract contour, and subsample to have a reasonable number of landmarks along the contour (reasonable meaning representative of the shape, but not to numerous to avoid overfitting and computational difficulties).
  – The shapes extracted must be visually checked, because the function may have run without error, but the contour may be badly represented. Here, data is lost by manually removing the badly extracted contours.
  – Automatically place landmarks, to allow the alignment and scaling of shapes and re-ordering of points. In this case, four landmarks could be automatically set on all shapes (Fig. 3). Landmark 1 (p1): point with largest averaged x,y value. Landmark 2 (p2): point with largest averaged x,y value on mirrored left side (using the absolute x value of all points with x < 0 after centering shape on its centroid). Landmark 3 (p3): point with smallest absolute difference from x=0 (after centering the shape on its centroid) in the lower half of the shape. Landmark 4 (p4): same as p3 for the upper half of the shape.
  – Shapes are then aligned and scaled using Procrustes analysis based on p1:p4.
  – Points of all shapes are re-ordered starting from p1, so that the ‘starting point’ of all contours is always the same, and the following points come in corresponding order, keeping the original direction of rotation.
- Apply Elliptical Fourier Analysis to the subsampled, aligned, scaled and reordered contour coordinates (Momocs function ‘efourier’).
- Apply Principal Component Analysis on the extracted Fourier coefficients.

**Figure 2:**
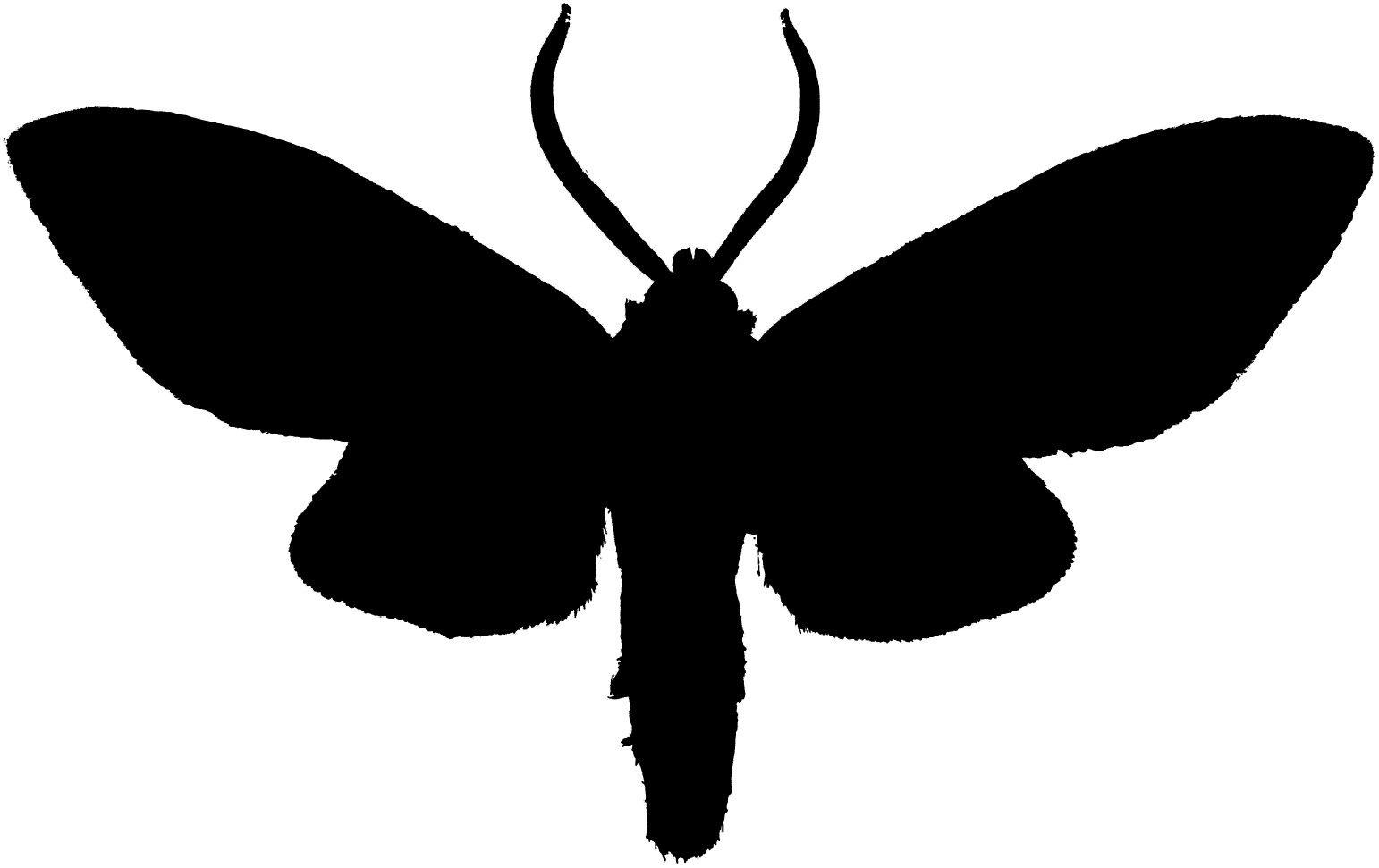
Example of binary (black/white) image obtained after putting the original images (e.g. Fig. 1) through the pipeline described here. The contour of the black shape can be automatically extracted using Momocs.

**Figure 3:**
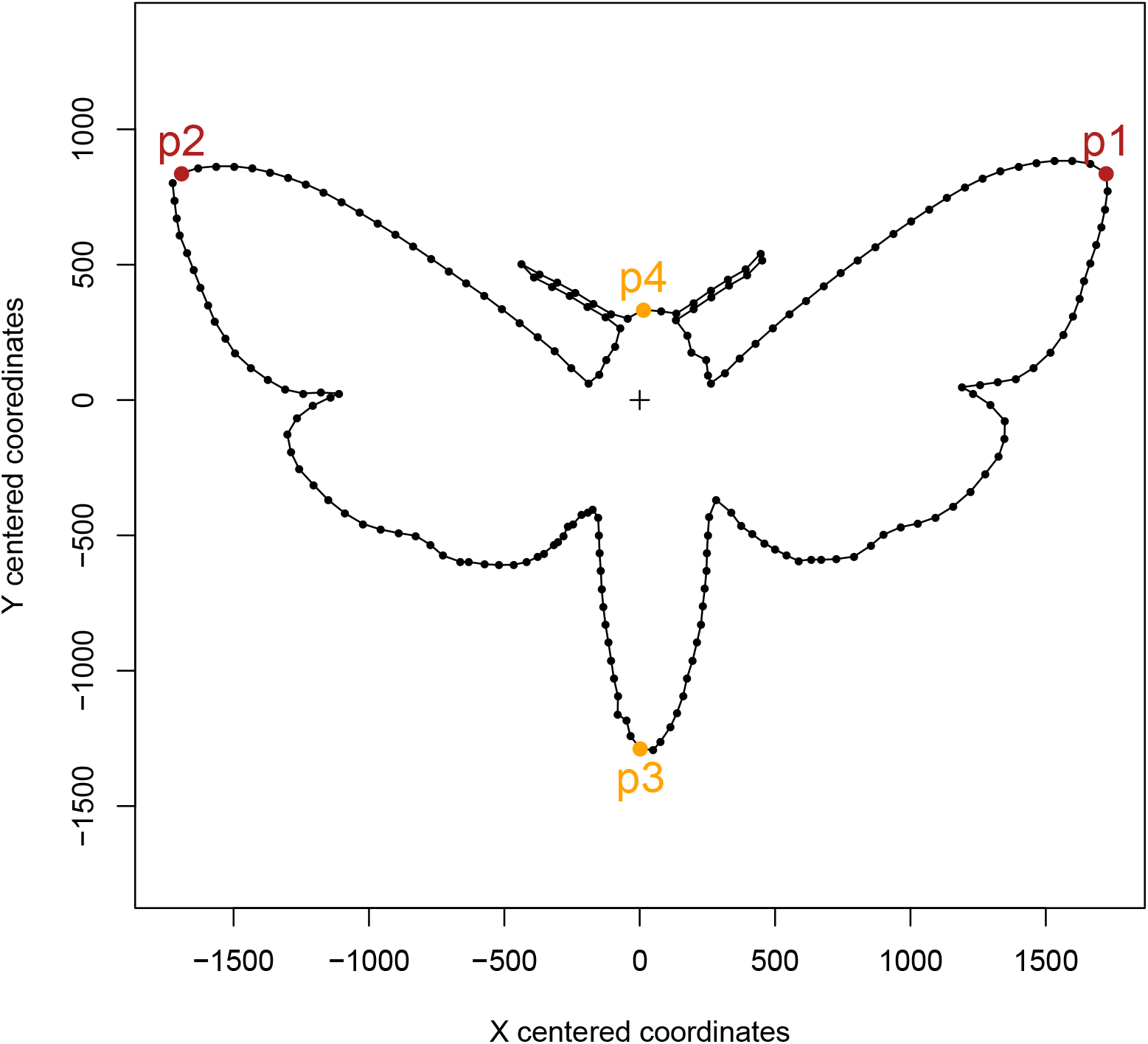
Example of extracted shape data. Yellow and red points labelled p1-p4 represent geometrically homologous landmarks that could be identified automatically on all shapes. The cross represents the centroid of the shape. These landmarks are used for alignment and scaling of the contours, and the ordering of the contour landmarks. Small black dots represent the contour landmarks after sub-sampling, which are then analysed using Fourier analysis.

Through this process, around 30% of the data were lost. Originally starting from 1991 (NHM) + 4516 (NHM Birdwing) + 1834 (MHNT) = 8341 individual specimen images, we managed to extract 1597 (NHM) + 2479 (NHM BirdWing) + 1715 (MHNT) = 5791 usable shapes. Although this loss is massive, the remaining dataset is nevertheless large, and the loss could probably be mitigated in the future by refining the pipeline. The remaining dataset was averaged per species for better readability of figures and to enable further analyses.

### 2.3 Validation of the automated data extraction pipeline

To validate the approach used in this article, we used a dataset of 2500 pictures of *Morpho* butterflies, obtained previously (Chazot et al., 2016), and for which morphometric data (landmarks and semi-landmarks) were obtained manually. The automated approach caused a loss of 50% of pictures, notably due to the fact that some pictures were very weakly contrasted (white butterflies on white background). The 1133 pictures from which contours were correctly extracted represented 22 species out of the 30 species of the original dataset. This dataset was then again restricted to specimens for which manually digitized landmarks were available for both the hindwing and forewing, and for which automatic contour extraction also worked correctly. Finally, for specimens for which dorsal and ventral views were available, only the contour extracted from the dorsal view was kept. After these restrictions, the final validation dataset contained 379 specimens, representing 19 species. Phylogenetic tree (pruned for the 19 species which were included in the validation dataset) and micro-habitat (canopy or understorey) data were also obtained from Chazot et al. (2016).

Three morphospaces at the individual level(i.e. PCAs based on landmark coordinates from each specimen) were produced: one for the forewing, one for the hindwing and one for the contour. The species-averages for the first 10 PCs were computed and plotted in these morphospaces, as well as the phylogenetic tree. To test whether phylogenetic and / or taxonomic signal was present in the automated contour morphospace, i) we computed Blomberg’s K for species average across each PC (function ‘phylosig’ from package phytools; Revell (2012)); and ii) using 1000 iterations of permutations between individuals, we randomized the species distributions along the first 10 PCs (representing >90% of total variance) of the automated contour morphospace, computed their randomized averages, and produced distributions of Procrustes distances between these random averages and the averages from the hindwing and forewing morphospaces. The true Procrustes distance between the species averages from the automated contour analysis was then compared to the null distributions produced, to test whether it was smaller than expected by chance.

### 2.4 Test of inter-museum bias in the main dataset

Because our main dataset comes from two different institutions (MHNT and NHM), technical differences in the preparation of specimens and/or digitization could produce systematic differences in the shapes, biasing the analysis. A direct way to test this would be to check whether matching species between datasets systematically differ. However, the datasets used here do not have such matching species. As an indirect test, we therefore studied two families that are common to the two collections, and include a sufficient number of species: i) the Nymphalidae, represented by 179 species in our MHNT dataset and 75 in our NHM dataset, and ii) the Papilionidae represented by 26 species from the MHNT and 27 from the NHM. The family averages and their associated 95% confidence intervals (CI) were computed, with the null expectation that the CI would overlap, showing no systematic difference between collections. However, an absence of overlap could suggest either systematic differences or a sampling effect.

## 3 Results/Discussion

### 3.1 Results for the validation dataset

The validation dataset used here first showed that our automated approach clearly fails with specimens that are not well contrasted from their background (e.g. light-colored butterflies over white background). After restricting the manually and automatically extracted datasets to common specimens, we were still able to produce morphospaces (Fig. 4) that included the majority of species (19 out of 30).

**Figure 4:**
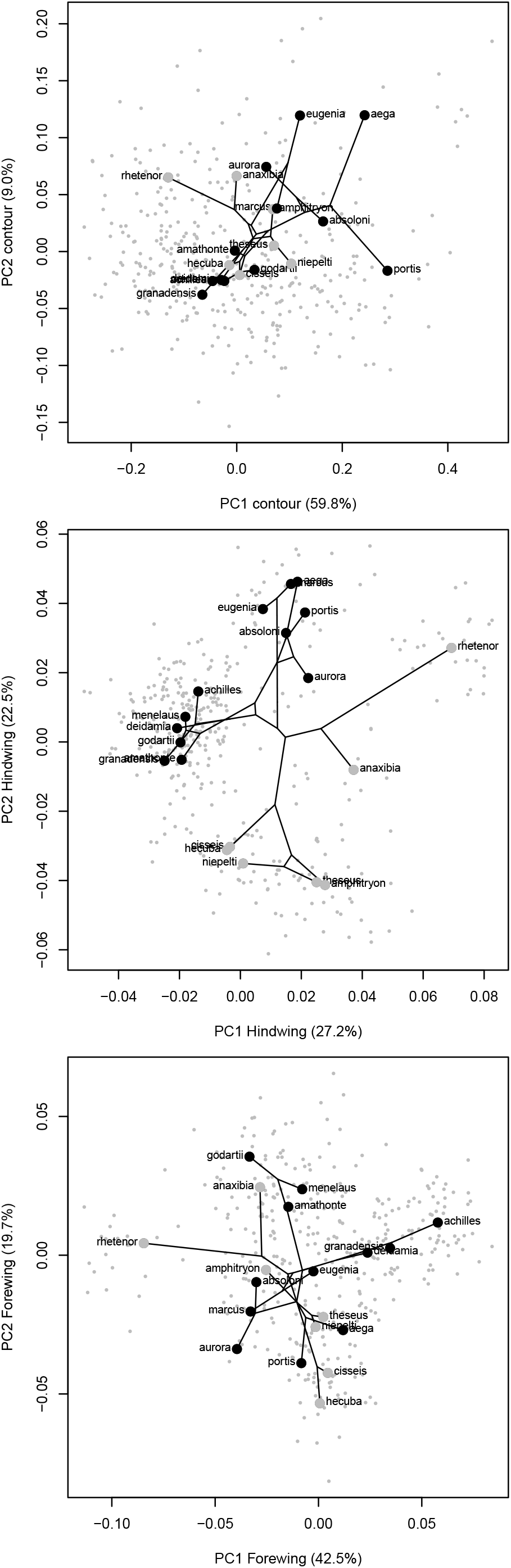
Morphospaces (PCAs) based on individual butterflies for which all three types of data were availables. Left: automatically extracted contour; middle: manually landmarked hindwing; right:manually land-marked forewing. Small grey dots represent individuals, while large dots represent species average (grey: canopy-dwelling species, black: understorey-dwelling species). Phylogenetic tree was mapped onto the graph using function ‘phylomorphospace’ from package ‘phytools’.

It is clear that the distribution of individuals in the contour morphospace (Fig. 4) is much less structured than in the manually landmarked morphospaces. In this regard, it is particularly noticeable that the PC1 of the contour morphospace gathers much more variation than the PC1 from the other morphospaces, suggesting it includes some amount of noisy variation. However, it also appears from the contour morphospace that sister species are often located close to each other. This is confirmed by significant phylogenetic signal along PC1 (K=0.86, P=0.001 based on 1000 iterations). In addition, our permutation test shows that the positions of species averages in the multivariate morphospaces globally (considering the first 10 PCs), are more similar than expected by chance alone (Fig. 5). On the other hand, the marked ecological differentiation between understorey and canopy species, which is observed in the hindwing morphospace, does not appear in the contour morphospace across the first 10 PCs (not shown in Fig. 4). In any case, differences between the morphospace were to be expected, considering that the shapes being used are widely different (whole contour including body and antennae vs. hindwing only vs. fore wing only). It is therefore in fact quite remarkable that our approach produces a space even fairly similar to the manual approaches.

**Figure 5:**
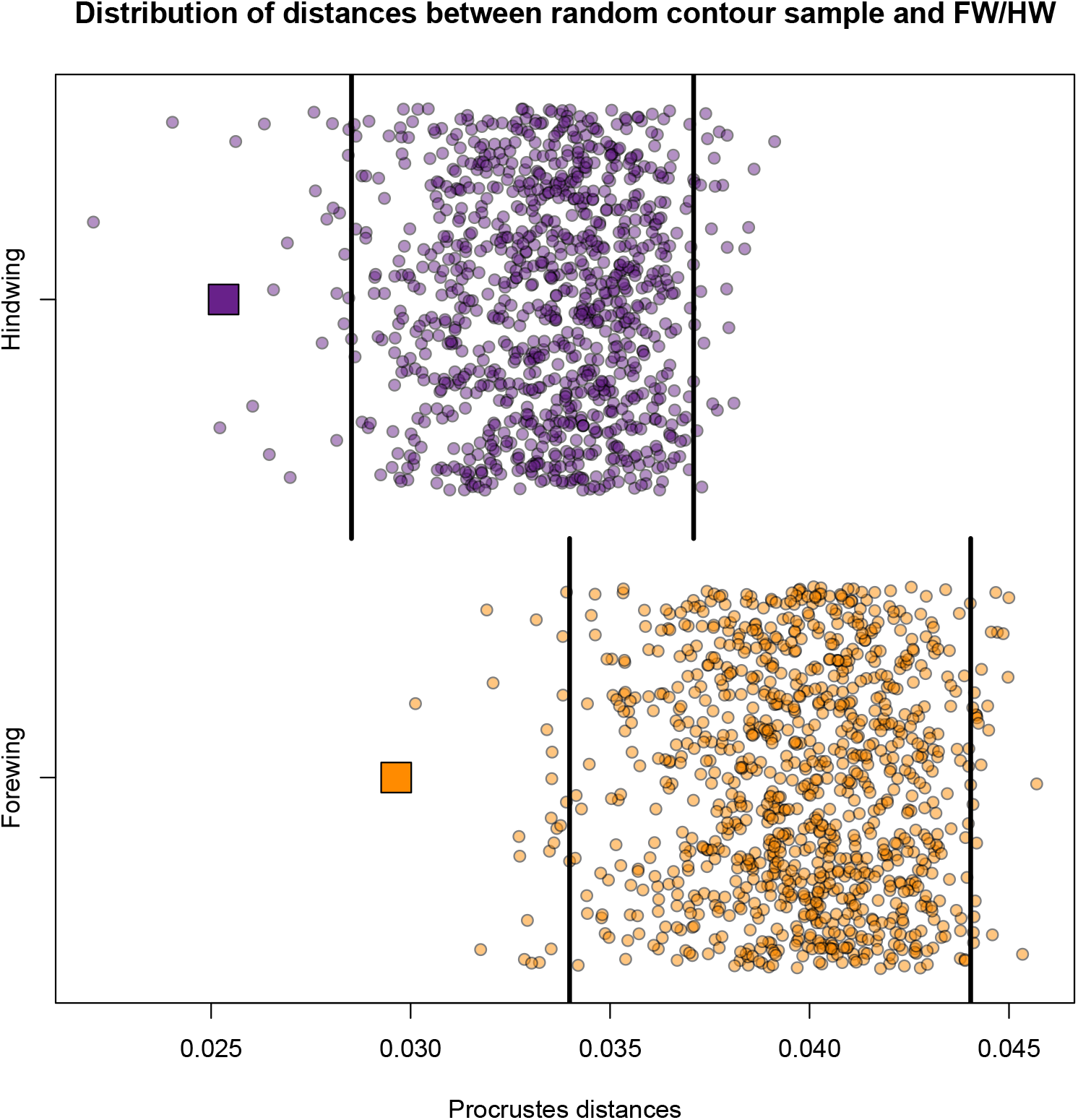
Procrustes distances between the species-averages of the contour morphospace and the species-averages of the hindwing morphospace (purple) and forewing morphospace (orange). Small circles represent the 1000 iterations of distances based on permutations of individuals in the contour morphospace. Large squares are the ‘true’ distances. Black vertical bars show the 5 and 95 percentile of the distance distribution generated by the iterations, showing that the true distances are smaller than expected by chance alone.

Although the matches between the morphospaces at the individual level, or even between the positions of the species averages, are not perfect, our results suggest that manually and automatically extracted data have some broadly common patterns which appear at the interspecific level. Differences may emerge due to more noise in the automatic extraction, but also due to the absence of certain species and to the fact that we use the contour which combines signals from both the fore and hindwings, as well as other parts of the body (notably the head/antennae which are not always in a standard position). Despite these limitations it still seems that biological signal is recovered from our data (here, phylogenetic and taxonomic). However, the morphological variation observed in this dataset may be relatively small, since it focuses only on the *Morpho* genus. Therefore the noise introduced by our method of data extraction may be very high in proportion to the morphological signal, leading to a fairly bad signal to noise ratio. Using the same approach in taxonomically wider datasets may leave the amount of noise unaffected, while increasing the morphological variation (i.e. biological signal), leading to an improvement of the signal to noise ratio. In any case, it would seem that the approach proposed here cannot be applied on datasets trying to document differences at scales smaller than between species of the same genus (i.e. individual level / intraspecific level). Improvements of the data extraction could be focused on the better standardisation of pictures, with better contrasts, but also on refining the parameters used for image segmentation.

### 3.2 Results for the dataset extracted from public online museum collections

Considering the previous results on the validation dataset, which included only one genus, we concatenated a new dataset based on semi-automatic extraction of contours from publicly available images of Lepidoptera collections from Natural History Museums. Although some data were lost in the process (again due to badly extracted contours notably related to contrast problems) we managed to semi-automatically produce a large dataset (over 5700 specimens) from public collections. The fact that some parameters must be set, and that the dataset must be manually checked for badly represented shapes limits the automation process. Still, the amount of time and effort needed for the whole process is much lower than for an equivalent dataset done manually, *even in the case where* collections and images would already be at hand. It is also notable that this type of automatically extracted data does not include subjective user bias as observed in manual landmarking, although other sources of error appear.

Figure 6 summarizes shape variation in our dataset at the interspecific level. The first two PCs of this morphospace represent about 75% of shape variation. The space is characterized by two areas of higher density, with one particularly obvious spot, in which Sphingidae species cluster. Shape variation along PC1, with over 57% of total variance is apparently driven by the width/length ratio, with more ‘rectangular’ or ‘triangular’ shapes (i.e. wide wings, short total length) along negative values and more ‘square’ shapes (i.e. width of wings about the same as total length) along positive values. Extreme positive values also include butterflies for which hindwings were wider than forewings, sometimes due to posterior extensions. On the other hand, variation along PC2 is harder to decript. It appears clearly that although Sphingidae is the family represented by the highest number of species, it also occupies a restricted area in comparison to other families, which produces the dense area visible on the negative side of PC1. It is also notable that some members of Nymphalidae and Erebidae seem to converge in terms of shape towards this Shpingidae pole. This was confirmed by a cross-validated LDA, which showed that all Sphingidae species were correctly assigned to their family, while 20% of Nymphalidae species were incorrectly assigned to Sphingidae and 10% to Lycaenidae). Among Lycaenidae, about 35% are incorrectly assigned as Nymphalidae. Among Erebidae (which are represented only by 14 species though), none were correctly assigned, with 7 going to Sphingidae, 6 to Nymphalidae and 1 to Papilionidae. This goes to show that taxonomic signal is not strong enough to allow perfect discrimination, aside from the very constrained Sphingidae, and that this constraint may be related to some functional variable, since at least two other families contain species that are close to the Sphingidae cluster. Especially conspicuous is the convergent morphology of some Erebidae species (*Himerarctia docis, Ernassa sanguinolenta, Mitochrista miniata*), less striking is the convergence of Nymphalids, probably just related to their developped forewing. *Zygaena transalpina* is also an interesting case of similar shape. Some ‘moths’ also show shapes similar to butterflies. Notably, they all show developped hindwings. One particularly impressive case is the Uraniidae.

**Figure 6:**
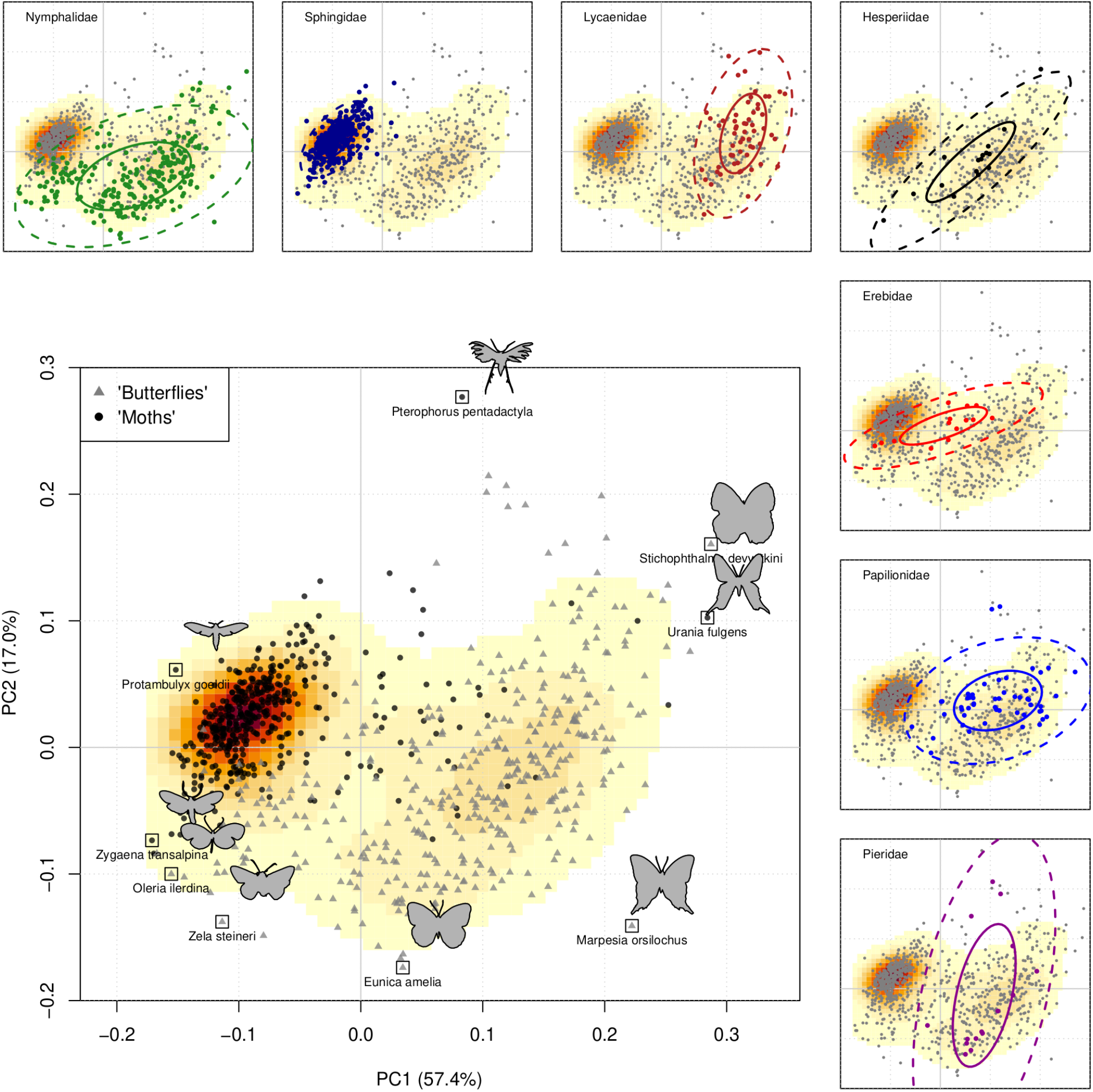
Species-level morphospace of 851 species of Lepidoptera. Each point represents one species average. The main panel includes contours of example individuals from the species forming the convex hull of the whole distribution over PC1 and 2. It also distinguishes between butterflies (grey triangles) and moths (black dots). Note that *Pterophorus pentadactyla* also has its legs digitized by our automated approach, which probably explains its outlying position. Background warm color gradient represents the density of points (warmer colors correspond to denser areas). Smaller panels around show the distributions of species for the families that included the largest number of species in our sample. Data ellipses include 50% (solid line) and 95% (dashed line) of species.

One difficulty with the previous interpretation of the morphospace is that all Sphingidae images come from the MHNT, making it difficult to disentangle the true morphological differences from differences that could be explained by each museum’s methods of preparation and digitization. To test this, we compared the distribution of Nymphalidae and Papilionidae species coming from both institutions. It appears from Fig. 7 that for both families, the distributions depending on the museums mostly overlap, suggesting that strictly museum related effects are not very strong and do not drive the distribution of species in this morphospace. The family per museum averages show no significant differences, except for PC2 in the Nymphalidae. Although this difference may only be due to different sampling of species in the two museums, it cannot be ruled out that it may also relate to inter-user biases. A non-significant difference, but in the same direction, is visible along PC2 also in Papilionidae. In any case, this potential inter-museum bias appears rather limited, and clearly does not drive the major structure of the morphospace, since no difference between museums is observed along PC1.

**Figure 7:**
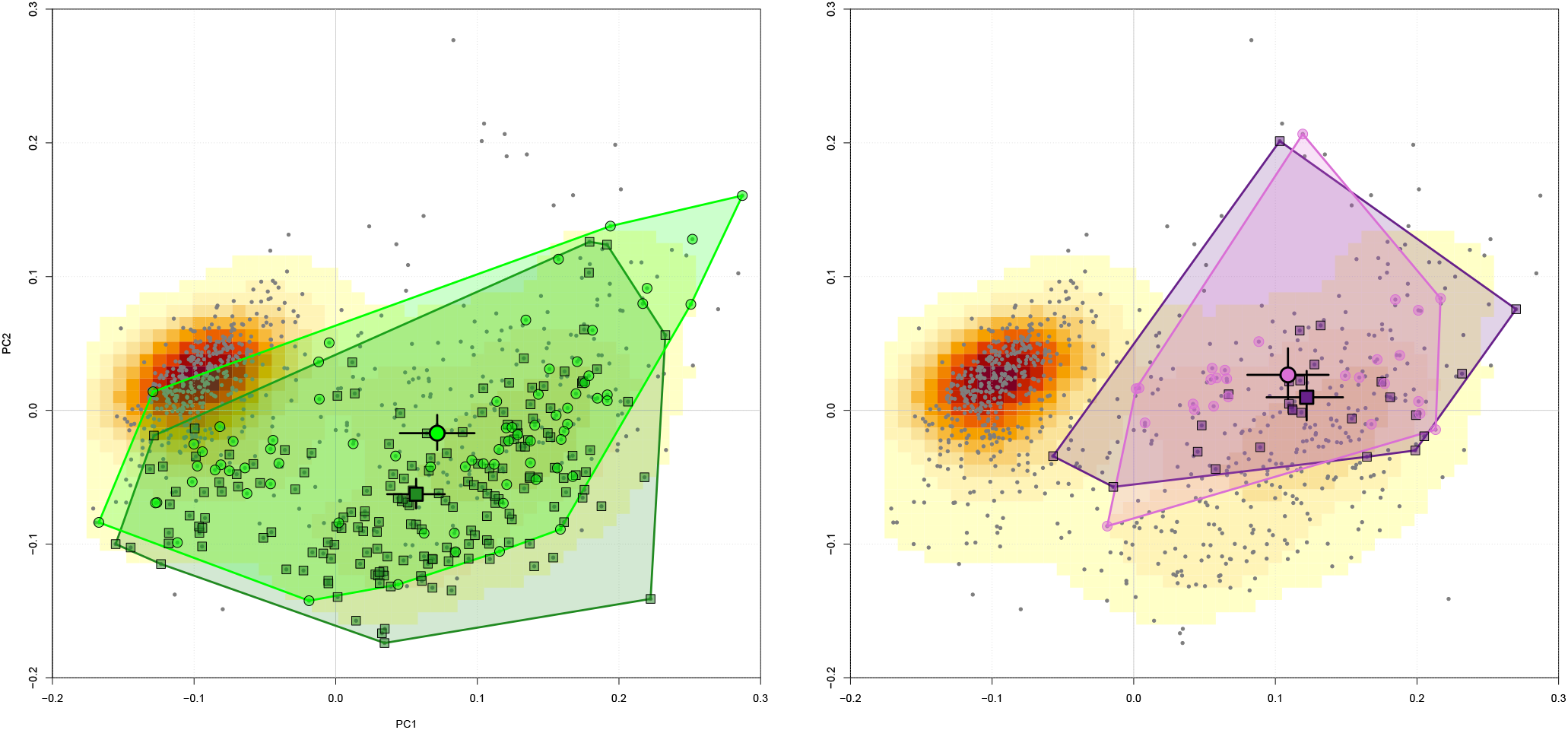
Same morphospace as in Fig. 6, highlighting Nymphalidae (left panel), and Papilionidae (right panel). Circles represent species from the NHM, while squares show species from the MHNT. Larger symbols represent the family average per museum, with black bars showing their associated 95% confidence intervals.

## 4 Conclusion

Houle et al. (2010) suggested that phenomics should increase both N (sample size) and P (dimensions), requiring as much as possible to automate the data collection. This is typically what we have done here, restricted to the morphological side of phenomics. However, going from our dataset, and averaging it to the species level could allow us to bind it with other phenotypic databases containing the same species. For example interesting ecological, functional, behavioural or life history traits to compare with morphological variation could be flight speed (maximum or average), flight mode (constant flapping, soaring, hovering…), migratory patterns or not, hovering flight or not, diurnal or nocturnal, etc. Since this type of data is becoming more and more available, morphometrics can follow this trend to add a new component of phenotype. Then again, basing our sampling on species which already have such data available is a good idea, but may not always be applicable.

Major limitations of this study are : 1) the lack of a phylogeny including the same species as we did in our main dataset. Solutions could be i) to include more species in phylogenetic studies, but this is probably not the easiest approach in super-diverse groups such as Lepidoptera; or ii) the other way around, to base the phenomic data collection on the sampling of already available phylogenies. 2) The absence of a size measurement, which is a major component of morphological variation and has allometric consequences on other components of morphology (i.e. the size, shape, form problem). Several possible solutions can be imagined: i) Do it manually based on measurements of scale bars, but this would defeat the purpose of automation, and may not be working in every case, if scale bars are absent. ii) Find available data on average size per species, but this would be restricted to the species level analysis and would require that this data is available for our sample. iii) Include a standard size measurement (e.g. body length) in the metadata associated with each specimen in collection, but this is a lot of work for museums, and has to be standardized strictly. iv) Automatize the process of scale bar measurement, which is challenging in terms of image treatment but is becoming tractable using machine learning but not applicable if scale bar is absent). 3) The automatic process described here is applicable only if the position of the biological object is fairly standard, and the superimposition of images requires that at least a few geometrically homologous landmarks can be identified automatically. Obviously there is probably no general easy way to automatize data collection over a range of very different objects, but if one is to restrict oneself to a certain group of organisms, the process described here can be fairly easy to adapt.

## References

V. Bonhomme, S. Picq, C. Gaucherel, and J. Claude. Momocs: Outline analysis using r. Journal of Statistical Software, 56(13):1–24, 2014. URL https://www.jstatsoft.org/v56/i13/.

A. Cardini, P. O’Higgins, and F. J. Rohlf. Seeing distinct groups where there are none: spurious patterns from between-group pca. Evolutionary Biology, 46(4):303–316, 2019.

N. Chazot, S. Panara, N. Zilbermann, P. Blandin, Y. Le Poul, R. Cornette, M. Elias, and V. Debat. Morpho morphometrics: shared ancestry and selection drive the evolution of wing size and shape in morpho butterflies. Evolution, 70(1):181–194, 2016.

R. Crowther, L. Allan, M. Barclay, and B. Huertas. Papilionoidea new types digitisation project [data set]. Natural History Museum, 2019a. doi: https://doi.org/10.5519/0015264.

R. Crowther, E. Devenish, P. Kokkine, N. Lowndes, P. Wing, L. Allan, J. Ayre, M. Barclay, and G. Martin. icollections: British and irish pyraloidea (moths) collection [data set]. Natural History Museum, 2019b. doi: https://doi.org/10.5519/0097198.

A. Goswami, A. Watanabe, R. N. Felice, C. Bardua, A.-C. Fabre, and P. D. Polly. High-density morphometric analysis of shape and integration: the good, the bad, and the not-really-a-problem. Integrative and Comparative Biology, 59(3):669–683, 2019.

D. Houle, D. R. Govindaraju, and S. Omholt. Phenomics: the next challenge. Nature reviews genetics, 11 (12):855–866, 2010.

C. Le Roy, V. Debat, and V. Llaurens. Adaptive evolution of butterfly wing shape: from morphology to behaviour. Biological Reviews, 94(4):1261–1281, 2019.

M. D. Lürig. phenopype: a phenotyping pipeline for python. Methods in Ecology and Evolution, 13(3): 569–576, 2022.

G. Pau, F. Fuchs, O. Sklyar, M. Boutros, and W. Huber. Ebimage—an r package for image processing with applications to cellular phenotypes. Bioinformatics, 26(7):979–981, 2010. doi: 10.1093/bioinformatics/btq046.

A. Porto and K. L. Voje. Ml-morph: A fast, accurate and general approach for automated detection and landmarking of biological structures in images. Methods in Ecology and Evolution, 11(4):500–512, 2020.

A. Porto, S. Rolfe, and A. M. Maga. Alpaca: A fast and accurate computer vision approach for automated landmarking of three-dimensional biological structures. Methods in Ecology and Evolution, 12(11):2129–2144, 2021.

L. J. Revell. phytools: An r package for phylogenetic comparative biology (and other things). Methods in Ecology and Evolution, 3:217–223, 2012.

R. J. Wilson, A. F. de Siqueira, S. J. Brooks, B. W. Price, L. M. Simon, S. J. van der Walt, and P. B. Fenberg. Applying computer vision to digitised natural history collections for climate change research: Temperature-size responses in british butterflies. Methods in Ecology and Evolution, 2022.

P. Wing, L. Allan, J. Ayre, M. Barclay, B. Huertas, L. Livermore, and G. Martin. Birdwing butterfly collection [data set]. Natural History Museum, 2019. doi: https://doi.org/10.5519/0014723.

